# Enhancer-gene regulatory interactions in colorectal cancer revealed through genome-wide CRISPRi perturbations

**DOI:** 10.64898/2026.06.05.730086

**Authors:** Philip J Law, Jayaram Vijayakrishnan, James Smith, Timothy Barry, Maria Mandelia, Eugene Katsevich, Richard S Houlston

**Author notes:** Correspondence: Richard Houlston. Nucleome Therapeutics, Oxford Science Park, Oxford, OX4 4GE, UK.

## Abstract

**Background:** Establishing functional relationships between distal enhancers and their target genes is a central challenge in genome biology and cancer genetics. CRISPR/dCas9-mediated perturbations followed by single-cell RNA sequencing (Perturb-seq) has emerged as an efficient means of establishing enhancer-target gene relationships.

**Results:** We used genome-wide CRISPRi Perturb-seq with 35,139 guide RNAs targeting 12,117 enhancers to create a functional enhancer-gene map of colorectal cancer (CRC), identifying 238 significant regulatory associations (FDR < 0.1). Integration of chromatin accessibility (ATAC-seq), histone modification profiling (ChIP-seq), and ultra-high resolution chromatin conformation capture (Micro-C) data revealed that enhancer regulation in CRC is constrained by topological domains and often targets the nearest gene, findings corroborated by Activity-by-Contact modelling.

**Conclusions:** We present here a comprehensive, genome-scale functional atlas of enhancer-gene associations in CRC. This resource should provide a foundation for studying gene regulation in colorectal tumorigenesis and for prioritising candidate non-coding drivers in cancer sequencing studies.

**PLAIN LANGUAGE SUMMARY:** *Mapping the hidden DNA switches that control colon cancer genes:* The total amount of DNA in humans is vast, but only a tiny proportion of it is directly related to genes. While people may know that genes act as blueprints for the body, there are also short stretches of DNA called “enhancers”, that fine-tune how strongly genes are activated. When these enhancers get damaged or mutated, it can trigger disease, suggesting that changes to these “control switches” themselves, and not just the genes, can be a driver of cancer. Thus, understanding how these enhancers function is key to uncovering the mechanisms that drive colorectal cancer. In this study, we aimed to map the role of enhancers in colorectal (bowel) cancer development. We used Perturb-seq, a cutting-edge genetic tool that allowed us to systematically turn off more than 12,000 individual enhancers, to identify which ones control which genes. We successfully linked 238 enhancers to the genes they regulate. Combining these results with other biological information, such as the 3D-shape of the DNA, allowed us to build a detailed picture of how enhancers physically interact with their target genes inside cancer cells. This work provides a comprehensive “wiring diagram” of enhancer-gene connections in colorectal cancer, and a resource for researchers seeking to develop new treatment approaches.

## INTRODUCTION

Enhancers are non-coding DNA elements that orchestrate cell-type-specific gene expression programs^1^. Through long-range interactions, they regulate transcriptional networks that underpin development, differentiation, and disease. Although epigenomic profiling, using ATAC-seq and histone ChiP-seq, has catalogued thousands of putative regulatory elements, biochemical signatures alone are insufficient to determine which enhancers are functional and which genes they regulate. A major unresolved problem is therefore linking candidate enhancers to their *bona fide* target genes^2^.

In cancer, dysregulation of enhancer activity can rewire transcriptional circuitry, promoting tumour initiation and progression^2^. While coding driver mutations have been extensively catalogued^3^, the non-coding genome remains comparatively unexplored. Statistical identification of non-coding drivers is hindered by the vast size of the search space and regional variation in background mutation rate. Tissue-specific regulatory elements represent a biologically enriched part of the genome in which to search for driver mutations, but such efforts depend critically on accurate enhancer-gene maps.

Multiplexed CRISPRi screens coupled with single-cell transcriptomics (Perturb-seq, also termed CROP-seq or crisprQTL) have recently emerged as a powerful approach for functional enhancer interrogation^4–9^. In Perturb-seq-style experiments, guide RNAs are introduced at high multiplicity of infection, enabling individual cells to harbour unique combinations of perturbations within an otherwise isogenic background. By assaying gene expression at single-cell resolution, it therefore becomes possible to infer regulatory relationships without requiring prior assumptions regarding enhancer targets.

To define the regulatory landscape of colorectal cancer (CRC), we performed genome-wide CRISPR/dCas9-ZIM3-KRAB perturbation of 12,117 candidate enhancer regions in the HT29 CRC cell line, followed by single-cell transcriptomic profiling. By integrating these data with chromatin accessibility, histone modification profiles, and Micro-C-defined three-dimensional genome architecture, we generated a high-resolution functional map of enhancer-gene interactions in CRC (**Fig. 1**). Beyond the specific associations identified, these data provide a framework for interpreting non-coding genetic variation and dissecting transcriptional circuitry in colorectal tumourigenesis.

**Figure 1:**
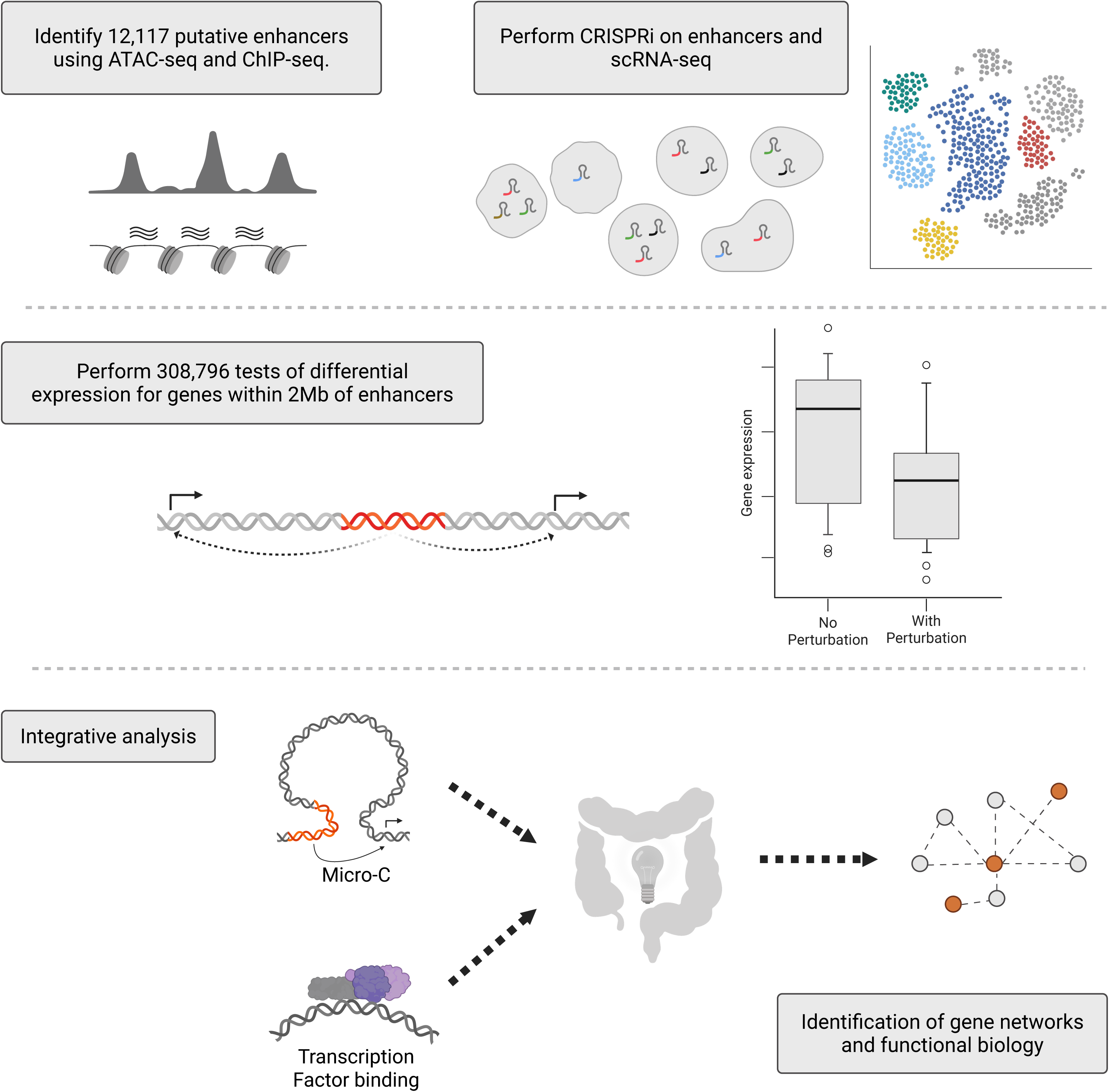
Experimental overview. Candidate enhancer regions were identified via ATAC-seq and H3K27ac and H3K4me1 ChIP-seq across multiple CRC cell-lines. Guide RNAs (gRNAs) targeting these regions were delivered via CRISPRi, followed by single cell RNA-seq. Differential gene expression was assessed for genes within 2Mb of each enhancer. Integration of Micro-C chromatin data and transcription factor binding, enabled identification of gene networks that potentially influence CRC development. Figure created using BioRender.

## RESULTS

### Genome-wide CRISPRi perturbation of candidate CRC enhancers

We identified 12,117 candidate enhancer regions in colorectal cancer (CRC) by intersecting ATAC-seq peaks with H3K27ac and H3K4me1 ChIP-seq marks across six CRC cell lines. Regions present in at least five of six lines were retained to enrich for shared, lineage-relevant regulatory elements. Up to three guide RNAs (gRNAs) were designed per enhancer, generating a library of 35,139 gRNAs cloned into a lentiviral CROP-seq vector^6^. The library was transduced at high multiplicity of infection (MOI) into HT29 cells expressing catalytically inactive Cas9 fused to the ZIM3-KRAB repressor domain (dCas9-ZIM3-KRAB), enabling CRISPR interference (CRISPRi)-mediated heterochromatin formation across an approximately 1-2 kb window surrounding each target site^10^. Ten days post-transduction, we profiled 252,485 single-cell transcriptomes. Across 61,541 detected genes, a mean of 57,459 reads per cell mapped to a median of 4,959 genes per cell. Sequencing metrics were robust, with Q30 scores exceeding 90% across barcodes and cDNA.

Conventional assignment of gRNAs to cells based on UMI count thresholding does not account for cell-to-cell heterogeneity in sequencing depth or batch effects^4^. We therefore used SCEPTRE^11^, which uses a mixture model^12^ to estimate the probability of gRNA presence in each cell and a conditional resampling approach for association testing. At a probability threshold of 0.8, the median MOI was 10 gRNAs per cell, and each gRNA was observed in a median of 67 cells (**Fig. 2A-D**, **Fig. 3**). Lowering the threshold to 0.5 did not significantly impact the estimated MOI estimates. gRNAs targeting the same enhancer were grouped using the “union” option in sceptre, prior to gRNA-to-gene association tests.

**Figure 2:**
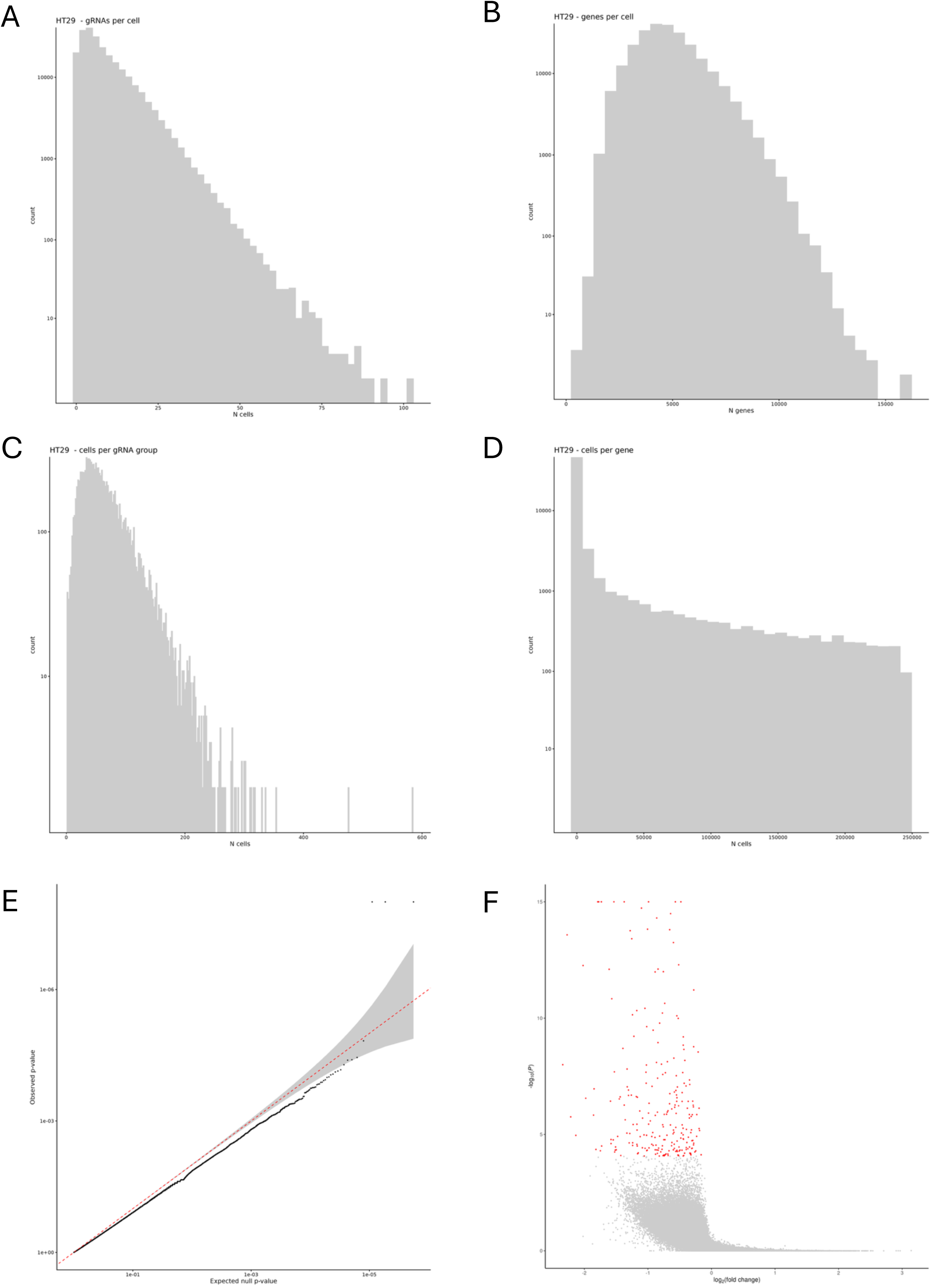
Summary of Perturb-seq data. Median gRNA per cell **(A)** and median genes per cell **(B)** were 10 and 4,628 respectively. Each gRNA was represented in a median of 57 cells **(C)** and each gene in 139 cells **(D)**. **(E)** QQ-plot of negative control region-gene-pairs shows well calibrated *P*-values. **(F)** Following QC, 278,667 enhancer-gene tests were performed within 2 Mb of enhancers, identifying 238 significant perturbations at *FDR* < 0.1 (red).

**Figure 3:**
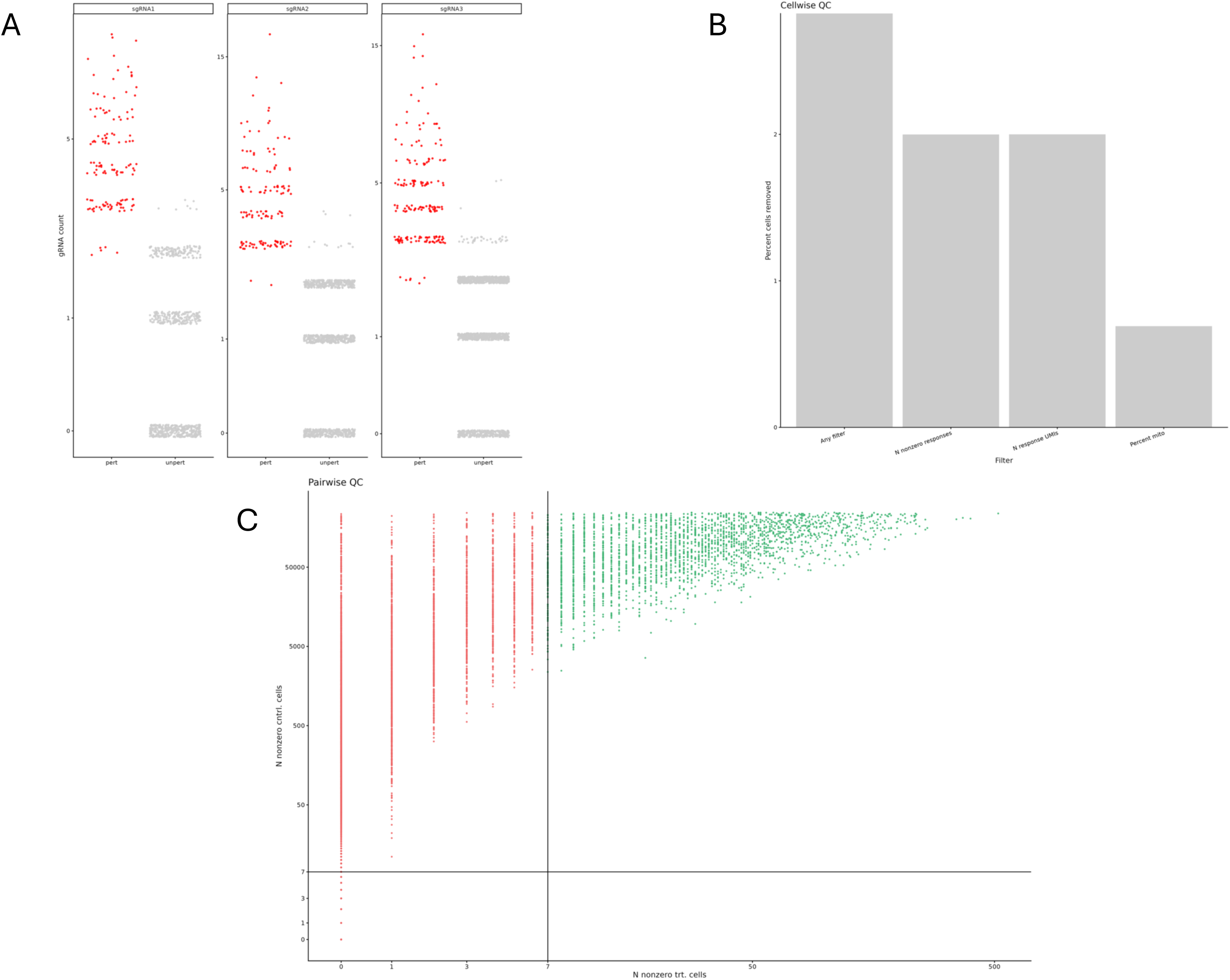
Details of the SCEPTRE QC analysis. **(A)** Visualisation of the assignment to cells of the three gRNAs mapping to the enhancer region chr1:184425472-184427021. The vertical axis represents the UMI count of the gRNA in a given cell. Cells containing the gRNA are classified as “perturbed” (pert, red) and those without as “unperturbed” (unpert, grey). **(B)** Cellwise QC statistics, showing the various QC filters applied to cells and the percentage of cells removed due application of a given QC filter. A cell may be flagged by multiple QC filters. **(C)** Pairwise QC was performed where each enhancer-gene pair was tested to ensure that there were sufficient gRNA reads to accurately identify a perturbation. “N nonzero trt. cells” represents the number of cells that contain the gRNA targeting the enhancer of interest, as well as having non-zero expression of the gene of interest. Conversely “N nonzero cntrl. cells” represents the number of cells that do not have gRNA targeting the enhancer and have non-zero expression of the gene of interest. Pairs failing the threshold (seven cells in each category) were excluded (indicated in red). 816,111 pairs were evaluated and 278,667 passed QC and were used in the final differential expression analysis.

We tested for differential expression of genes within 2Mb of the enhancer region, using sceptre’s conditional resampling-based approach^11^, performing 278,667 pairwise tests (median of 18 genes per enhancer). A matched analysis of negative control region-gene pairs indicated adequate control of false positive discoveries (**Fig. 2E**). Imposing a false discovery rate (FDR) threshold of 0.1 we identified 238 significant associations (**Fig. 2F**, **Table 1**).

**Table 1:** The 238 significant (FDR < 0.1) results from the SCEPTRE analysis annotated with functional data. “N_nonzero_trt” and “n_nonzero_cntrl” indicate the number of cells that express the gene, with and without the gRNA, respectively. “ABC Score” provides the score from the ABC analysis determining if the enhancer is affecting expression of the gene. The default threshold of 0.025 was used. “Micro-C” indicates if the enhancer region has a 3D contact with the TSS of the target gene. “Intronic” indicates whether the enhancer falls within the intron of the target gene. “Essential gene” indicates if the gene was determined to be essential in the DepMap CRISPR knockdown screens - HT29: essential in the HT29 cell line; Primary only: only primary CRC cell lines; Metastasis only: only in metastatic CRC cell lines; Primary&Metastasis: essential in both primary and metastatic CRC cell lines. The “Fitness score” columns refer to the target priority scores from DepMap, either in the CRC cell lines, or across all cancers (pancancer). A score of 40 is indicative of a potential therapeutic target.

### Architectural constraints shape enhancer-gene interactions

The identified significant associations revealed strong spatial constraints. Most associations occurred over relatively short genomic distances, with nearly two-thirds (64%; 154/238) involving genes within 100kb of the perturbed enhancer, and only a small faction extending beyond 1Mb (3%; 6/238). The majority of associations were confined within a single topologically associating domain (TAD) (83%; 197/238), which is consistent with three-dimensional genome organisation restricting regulatory communication. In more than half of the functional enhancers localised within introns of respective target genes (58%; 138/238), and in many cases, the enhancer targeted the nearest gene TSS (33%; 79/238). Although nearly all enhancers regulated a single gene, a subset (n=15) showed evidence of regulatory convergence, with multiple enhancers influencing the same transcript. Genes subject to multi-enhancer regulation included *CFTR* and *KITLG*, which are known to participate in cancer-relevant pathways^13,14^. Conversely, a small number of enhancers (n=7) affected the expression of multiple genes, including chr17:58390921-58392964 affecting both *RNF43* and *SUPT4H1*. A notable proportion of enhancer targets (29%; 70/238) were over-expressed in HT29 relative to normal colonic epithelial cells (>1.5-fold log₂ fold difference; **Fig. 4**), providing evidence of linkage of functional enhancer activity to oncogenic transcriptional programs.

**Figure 4:**
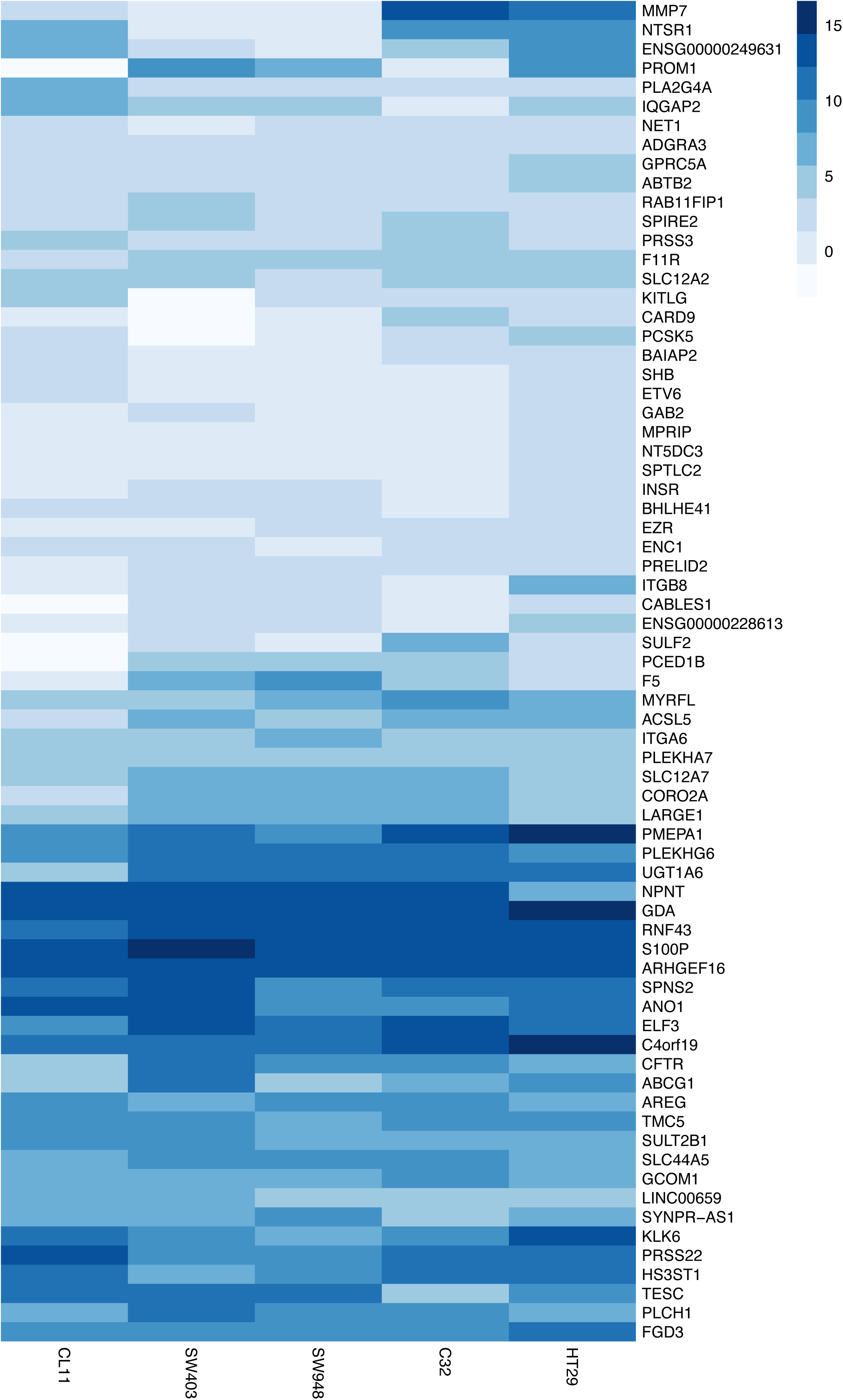
Heatmap of genes that are over-expressed (>1.5 log_2_-fold-change) in HT29 in relation to normal colon cells (HCEC-1CT). Also shown are the log_2_-fold-change for these genes in the C32, CL11, SW403 and SW948 CRC cell lines.

### Integration with chromatin structure and predictive modelling

To assess the relationship between physical chromatin architecture and functional regulation, we applied the Activity-by-Contact (ABC) model^15,16^ integrating ATAC-seq, H3K27ac ChIP-seq, RNA-seq, and Micro-C data in HT29. The ABC predictions provided support for 89% of significant Perturb-seq associations (ABC score > 0.025, **Table 1**), offering orthogonal validation of functional associations. Comparison with enhancer-gene predictions derived from other cell types highlighted pronounced tissue specificity. In a contemporaneous ENCODE study^17^ using Hi-C and ABC-modelling, 57% (136/238) of the predicted enhancer-gene associations were confirmed, albeit with many being predicted at low confidence. In contrast only 15% and 10% enhancer-gene associations were identified in lymphoblastoid cell lines GM12878 and K562, respectively.

### Regulatory networks and cancer relevance

Several enhancer-gene associations from our Perturb-seq screen have biological significance to the development of CRC. Repressing the enhancer region at chr20:47809086-47810869 was associated with reduced expression of *SULF2* (sceptre log_2_ fold-change (logFC) = -1.77, *P_adj_* = 2.93 x 10^-38^; **Fig. 5A**). The enhancer featured a Micro-C interaction with the transcription start site (TSS) of *SULF2* and is supported by the ABC modelling. Over-expression of *SULF2* has been associated with increased cell viability, proliferation, and migration, and poorer prognosis in CRC^18,19^.

**Figure 5:**
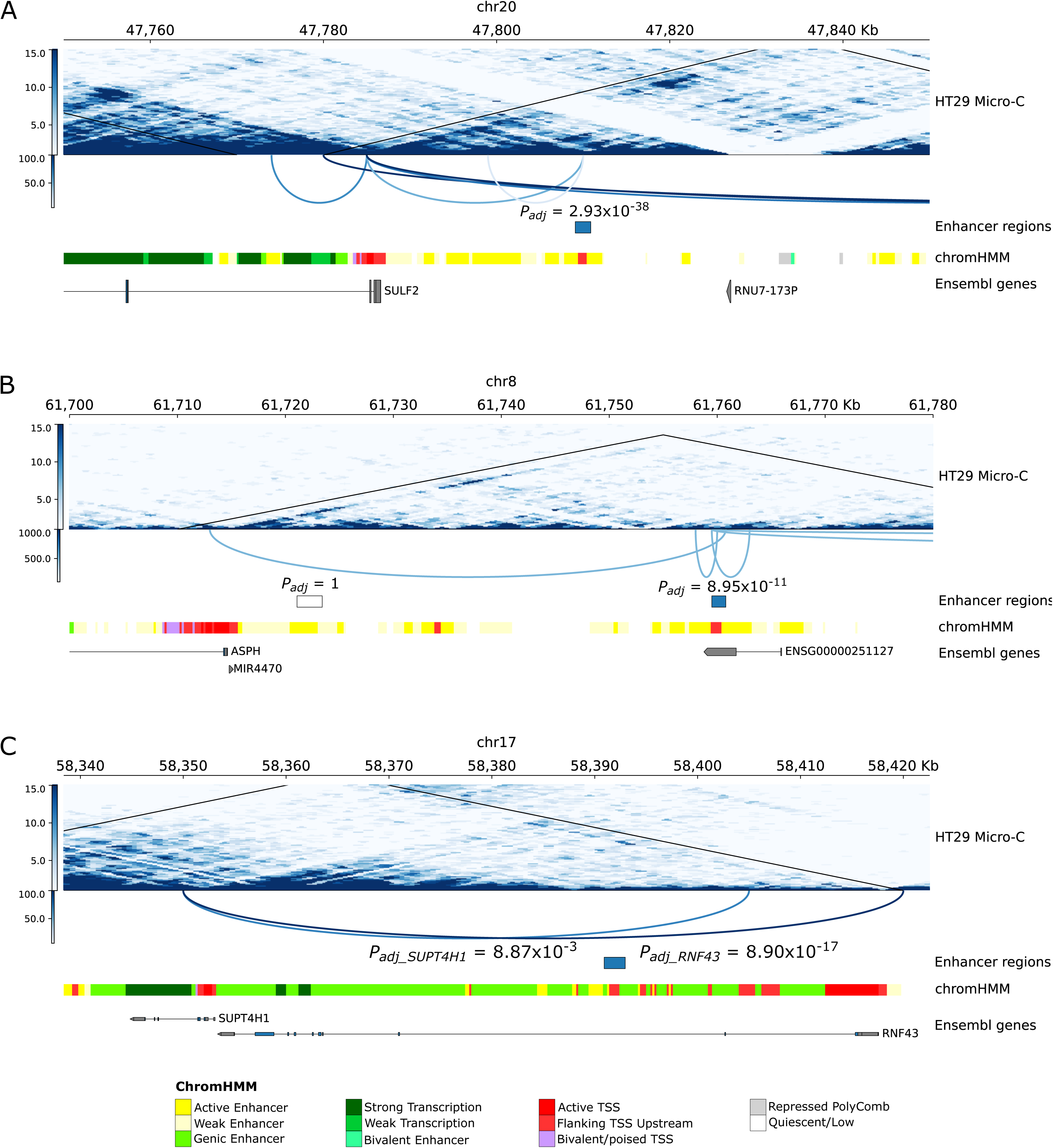
Locus plot of significantly perturbed regions. HT29 Micro-C contact matrices with predicted TAD boundaries (black lines) and significant enhancer interactions are shown for **(A)** *SULF2*; **(B)** *ASPH*; **(C)** *SUPT4H1* and *RNF43*. Arcs intensity reflects the interaction score from pyHICCUPS. ChromHMM enhancer annotations and Ensembl gene models are shown below. Enhancer *P*-values are from sceptre after multiple hypothesis testing correction; coordinates are in GRCh38. Figures generated with pyGenomeTracks^52^.

Similarly, the enhancer at chr8:61759498-61760795 implicates *ASPH* (logFC = -0.63, *P_adj_* = 8.95 x 10^-11^; **Fig. 5B**). Although another enhancer exists in closer proximity to the *ASPH*, this enhancer was not significant in the SCEPTRE analysis. *ASPH* is overexpressed in several cancers^20,21^ and is associated with increased migration and invasion of tumour cells^22^. Higher *ASPH* levels have been shown to be associated with lower rates of CRC survival^23^.

The enhancer at chr17:58390921-58392964 is an example of an enhancer impacting the expression of a multiple genes, targeting both *RNF43* (logFC = -0.57, *P_adj_* = 8.90 x 10^-17^) and *SUPT4H1* (logFC = -0.65, *P_adj_* = 8.87 x 10^-3^; **Fig. 5C**). *RNF43* is a negative regulator of Wnt-signalling^24,25^ and *SUPT4H1* is a determinant of CRC germline predisposition^26^.

While most enhancer-gene associations operated over short distances a notable example of a long-range association pairing at over 1.5Mb apart was between the enhancer at chr19:6863405-6863793 targeting the gene encoding the Ras-related protein Rab-11B (*RAB11B*; logFC = -1.98, *P_adj_* = 9.63 x 10^-4^).

Functional perturbations implicated numerous genes with established roles in colorectal tumorigenesis. Multiple myosin genes (*MYO10*, *MYO1E*, *MYO6*) were also identified, suggesting coordinated regulation of cytoskeletal and migratory programs. Transcription factor (TF) binding at functional enhancers using the JASPAR database frequently implicated KLF5, CTCF, and ZNF263, inferring the presence of structured regulatory networks. Focusing on genes over-expressed in HT29 (>1.5-fold vs normal colon; **Fig. 4**) and CRC-relevant TFs (e.g. *ETS2*, *SMAD3*) we defined major CRC regulatory networks: SP1-mediated activation, and ZNF263-mediatated repression (**Fig. 6**). Several genes associated with perturbed enhancers encode TFs that additionally have binding sites on multiple enhancer regions, including BACH1, NFIC and PPARD.

**Figure 6:**
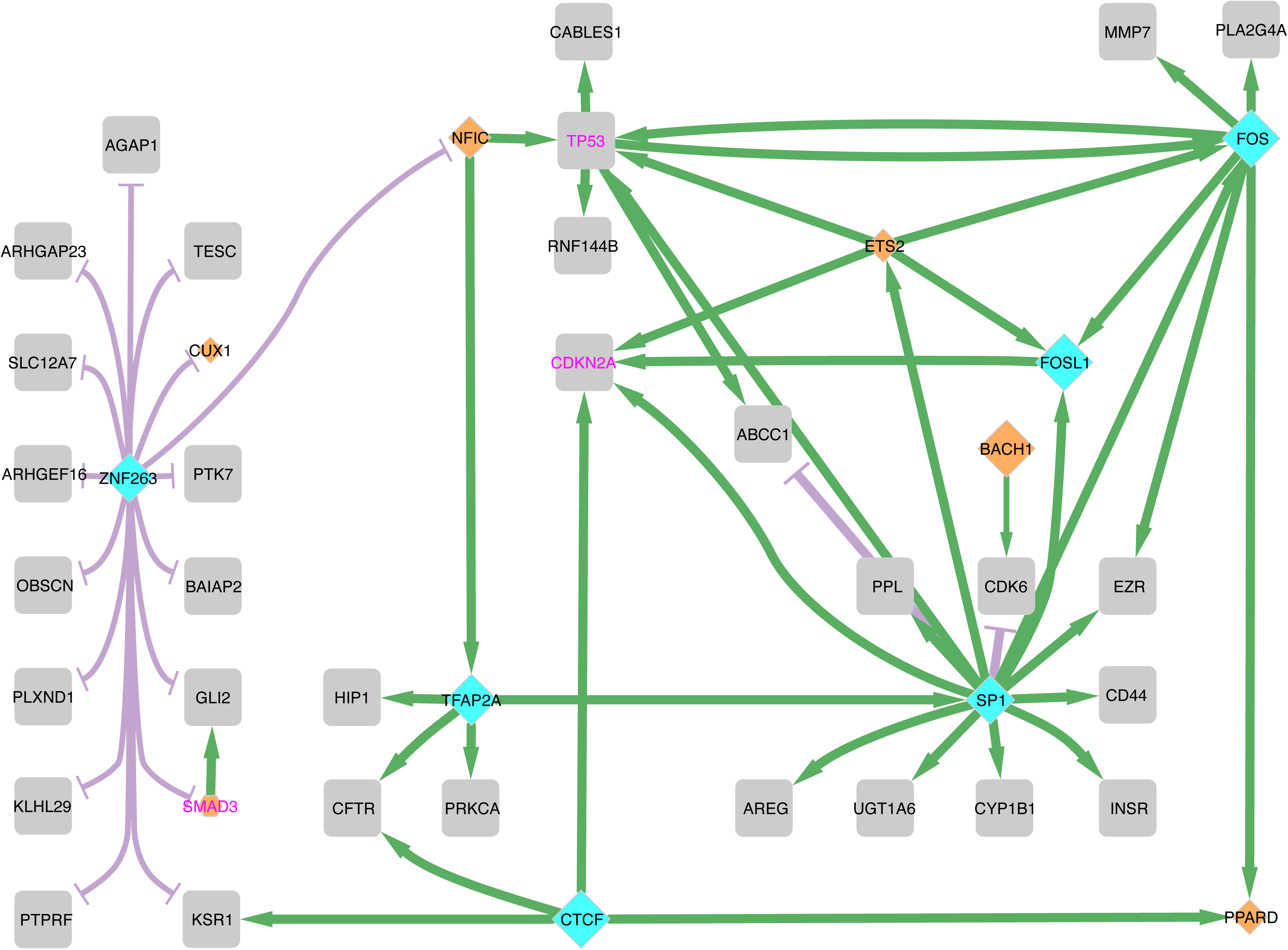
Network of perturbed genes and transcription factors. Genes overexpressed in HT29 and identified in Perturb-seq are connected based on high-confidence interactions (A/B scores). Green edges indicate activation; red edges indicate repression; edge thickness reflects interaction confidence. Predicted TFs bound to enhancer regions are shown in blue diamonds; TFs encoded by enhancer-perturbed genes are orange. Diamond size reflects frequency of binding site detection (30-115). Gene names in purple denote those mutated in CRC WGS studies.

Perturbed targets included recognised CRC coding drivers^3^ (*SMAD3*, *RNF43*, *EHD2*, and *ELF3*) as well as those previously identified through genome-wide association studies^27,28^ (GWAS) (*ANO1*, *SMAD3*, *STK39*, *PLEKHG6*, *GNA12*). Interestingly, the enhancer at chr11:69947520-69952396 (11q13.3), which is predicted to affect the expression of *ANO1* (logFC = -1.01, *P_adj_* = 3.39 x 10^-10^; **Fig. 7A**), does not overlap with the region identified in the GWAS. However, there is a Micro-C interaction between the enhancer region and GWAS region. Intriguingly, there are multiple transcripts of *ANO1*, with the shorter transcripts in proximity to the GWAS region, and the longer transcript in proximity to the enhancer region. The enhancer at chr15:67149995-67151277 was associated with perturbed expression of *SMAD3* (logFC = - 0.46, *P_adj_* = 6.23 x 10^-4^; **Fig 7B**), a key component of TGF-β signalling. Furthermore, several genes implicated by GWAS but were not perturbed (*MYC*, *CDKN1A*) still mapped to these networks, suggesting mechanistic overlap (**Fig. 8**).

**Figure 7:**
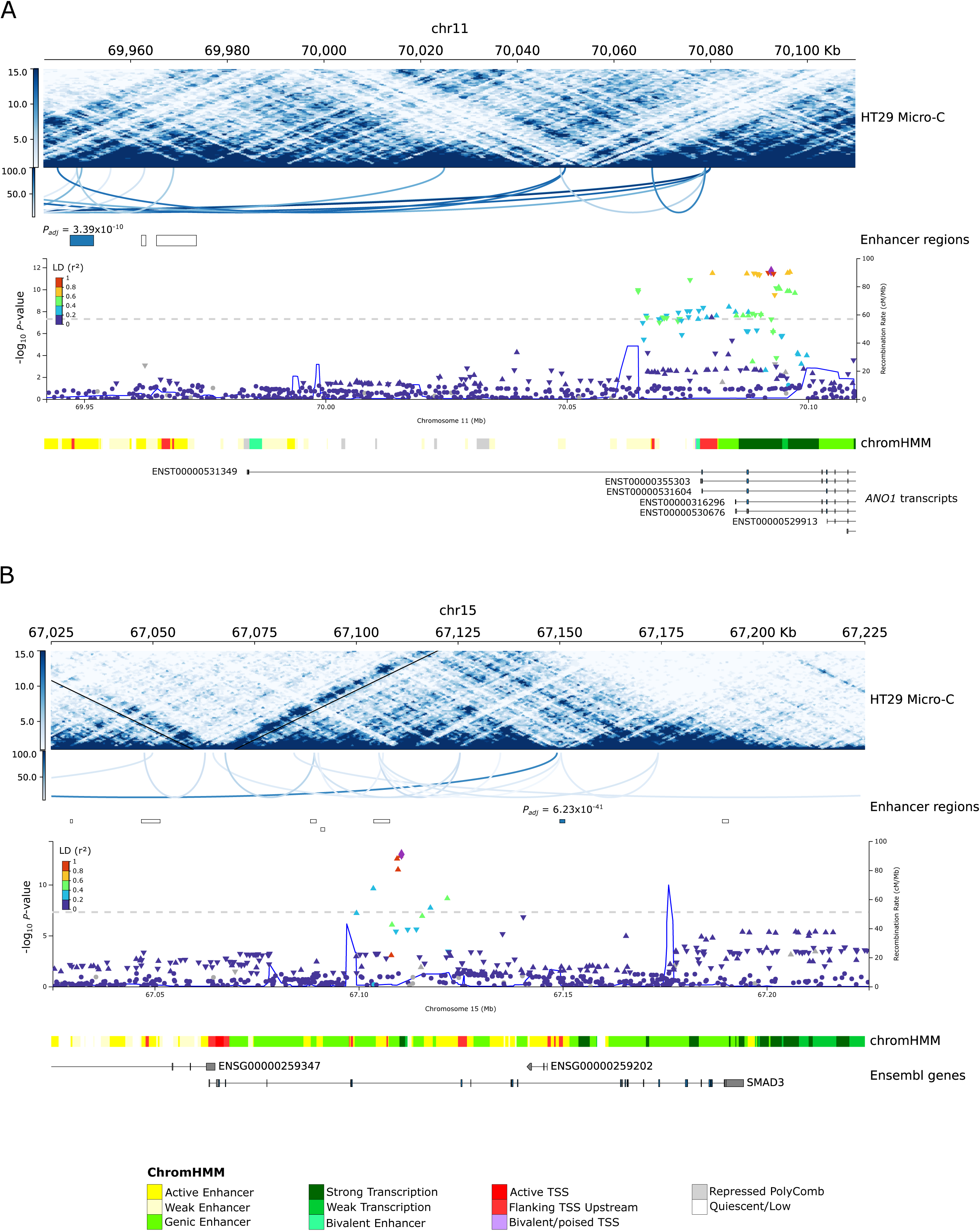
Locus plot of significantly perturbed regions with associations near known GWAS regions. (A) *ANO1*; (B) *SMAD3*. Shown are the HT29 Micro-C map together with predicted TAD boundaries (black lines), as well as significant interactions. Colour intensity is reflective of the interaction score from pyHICCUPS. Below are the Manhattans plots from the latest CRC GWAS^28^, showing each variants’ association with CRC (-log_10_*P*-value). At the bottom are the predicted annotation from chromHMM based on in-house histone data, and the gene models from Ensembl. Enhancer *P*-values are from the sceptre analysis after multiple hypothesis testing correction. Any unannotated enhancers have *P_adj_*= 1. All coordinates are in GRCh38. Figures generated using pyGenomeTracks and LocusZoom.

**Figure 8:**
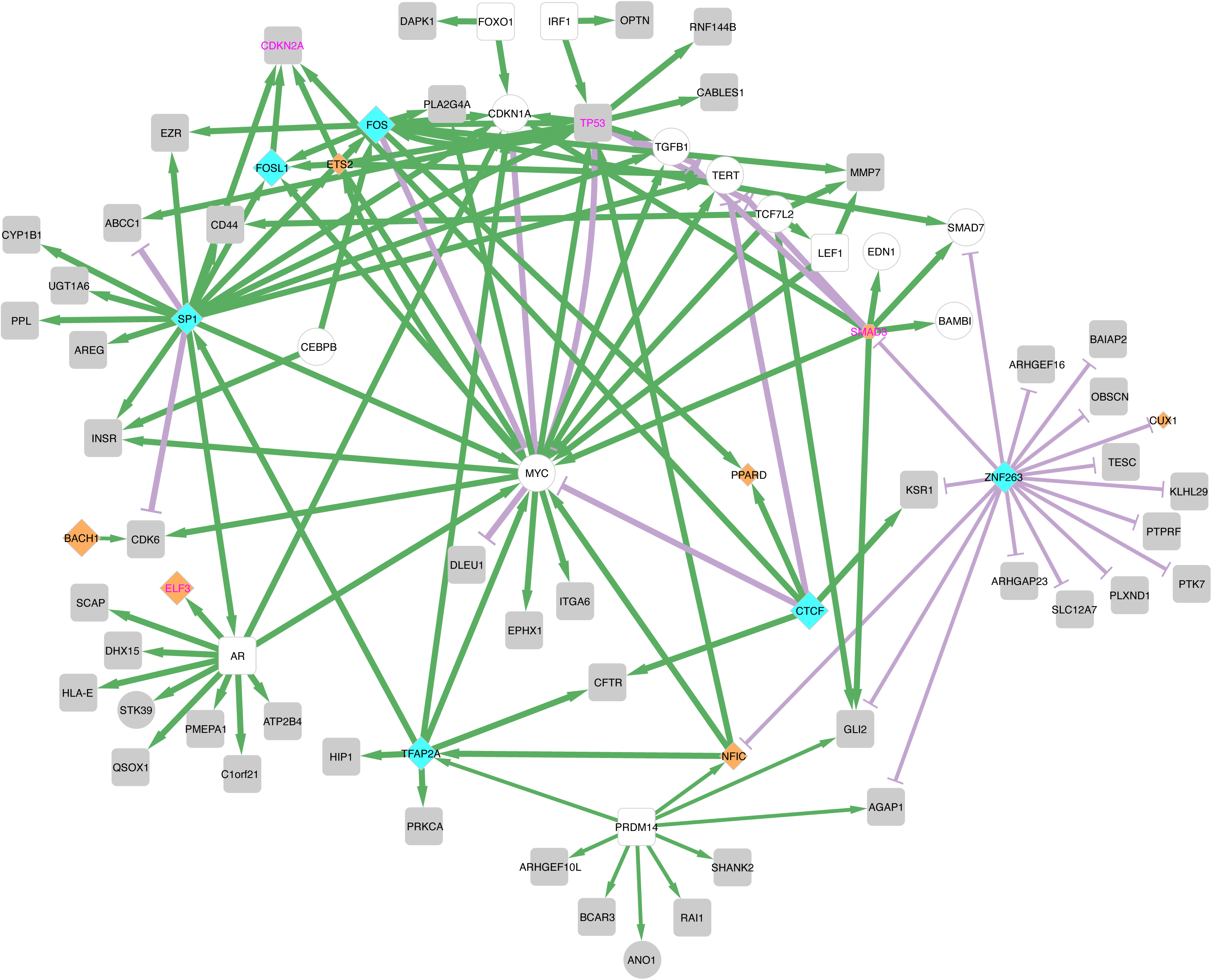
Expanded version of the regulatory network figure. Including genes implicated through GWAS (round shapes), highlighted additional connections. Genes in white shapes were not found through the perturbSeq analysis, but regulate genes that were. Green edges indicate activation; red edges indicate repression; edge thickness reflects interaction confidence. Predicted TFs bound to enhancer regions are shown in blue diamonds; TFs encoded by enhancer-perturbed genes are orange. Diamond size reflects frequency of binding site detection (30-115). Gene names in purple denote those mutated in CRC WGS.

Interrogation of CRISPR knockout dependency data from DepMap identified a subset of 46 enhancer-regulated genes (**Table 1**), including *SCAP*, *CDK6* and *LUC7L3* that are essential for tumour fitness. These findings connect distant regulatory elements to genes with potential therapeutic relevance and suggest that enhancer dysregulation may contribute directly to cancer cell dependencies.

## DISCUSSION

The challenge in deciphering the regulatory landscape of cancer genomes is identifying and validating the genes that are the target of potential enhancers. By combining genome-scale CRISPRi with single-cell transcriptomics and integrated multi-omic modelling, we have generated a functional evidence-based enhancer-gene map in CRC.

Three principal insights emerge from our analysis. First, enhancer activity is strongly constrained by three-dimensional genome architecture. Most functional interactions occurred within the same TAD, involved short genomic distances, and targeted the nearest gene. In predictive modelling, shared TAD membership, chromatin contact frequency, linear proximity, and the number of intervening genes each contributed modestly, indicating that enhancer-gene communication reflects a multifactorial integration of spatial organisation and chromatin context rather than a single, dominant determinant. Second, regulatory wiring is highly tissue specific. Concordance between functional perturbation and Activity-By-Contact based predictions was substantially higher in HT29 than in unrelated cell types, underscoring the limitations of transferring enhancer-gene assignments across lineages. Functional interrogation in disease-relevant cellular contexts is therefore essential for the accurate interpretation of non-coding elements. Third, the identified enhancer-gene relationships converge on genes central to colorectal tumorigenesis. Targets include established coding drivers, genes at GWAS loci, and genes essential for tumour cell fitness. Transcription factor motif enrichment further supports the existence of structured oncogenic and tumour-suppressive regulatory circuits embedded within the enhancer landscape.

From a cancer genomics perspective, this resource provides a functional scaffold for interpreting non-coding alterations. Restricting analysis to enhancers with experimentally validated targets improves prioritisation of recurrent point mutations and clarifies the consequences of structural variants, focal amplifications, and putative enhancer hijacking events. By anchoring non-coding lesions to defined gene targets, the map strengthens mechanistic inference and reduces ambiguity in driver assignment.

We acknowledge that the present analysis has limitations. Firstly, we have used a cellular model to define enhancer-gene relationships, which is unlikely to fully recapitulate CRC. Additionally, the enormous multiple testing burden incurred by conducting a genome-scale Perturb-seq screen has impacted on our power to identify many enhancer-gene pairings. Furthermore, biochemical effects may influence the detection of the perturbation, including incomplete or ineffective repression by the KRAB domain, enhancers only required for the initiation of gene expression, and the histone marks used here to define candidate enhancers excluding some classes of regulation^4^. Finally, there may be technical challenges. Large-scale screens require substantial sequencing depth and library complexity, and single-cell RNA sequencing data are inherently sparse, reducing sensitivity to detect modest expression changes. Despite these challenges, CRISPRi-based perturbation remains uniquely suited for unbiased high-throughput enhancer-gene mapping. Accepting these caveats, our analysis provides insights into key properties of enhancer-gene relationships and define regulatory gene networks in CRC.

## CONCLUSIONS

Through experimentally linked enhancer-gene regions, we provide a functional understanding of the regulatory mechanisms that control the gene expression underpinning CRC biology. These findings position enhancer regions as a biologically enriched and mechanistically interpretable compartment of the cancer genome for driver discovery. The resource generated here provides a foundation for integrating regulatory genomics with functional assays, advancing non-coding variant interpretation, biomarker development, and therapeutic prioritisation.

## METHODS

### Cell culture

HT29 (ACC299, DSMZ) cells were cultured in McCoy’s 5A (Modified) medium with GlutaMAX™ Supplement (36600021, Gibco). HEK239T (CRL-11268, ATCC), SW403 (ACC294, DSMZ), SW480 (ACC313, DSMZ) and SW948 (91030714, ECACC) cells were cultured in Dulbecco’s Modified Eagle Medium (61965026, Gibco). CL11 (ACC467, DSMZ) were grown in DMEM/F12 (Gibco), and C32 (ECACC) in IMDM (Gibco). The normal colon crypt cell line HCEC-1CT (CkHT039-0229, Evercyte) was cultured in 3% O_2_, 5% CO_2_ at 37°C in DMEM and M199 (Gibco) mixed in 4:1, supplemented with the ColoUp kit (Evercyte). All media was supplemented with 10% heat inactivated foetal bovine serum (FBS; F7524, Sigma), except CLL11 cells (20% FBS). Cells were cultured in a humidified incubator at 37°C in 5% CO_2_ and passaged at ∼90% confluence using TrypLE (1260402, Gibco).

### ChIPmentation

ChIPmentation for H3K4me1 (Diagenode, C15410194), H3K4me3 (Diagenode, C15410003), and H3K27ac (Diagenode, C15410210) was performed as described^29^. Briefly, 3-4 x 10^7^ cells were fixed in 1% methanol-free formaldehyde (10 min) and quenched with 200mM glycine, washed in PBS-A, flash-frozen in liquid nitrogen, and stored at -80°C.

For chromatin preparation, pellets were thawed on ice and lysed in ice-cold cytoplasmic lysis buffer (50mM HEPES, 150mM NaCl, 1mM EDTA, 1% Triton X-100, 0.1% sodium deoxycholate, 0.1% SDS) supplemented with protease inhibitors. Following centrifugation (1,200rpm, 4°C) pellets were incubated in a high-SDS lysis buffer (SDS concentration increased to 1%) and resuspended in sonication buffer (10mM Tris, 1mM EDTA, 0.1% SDS). Chromatin was sheared using a Covaris E220 (1ml Covaris Millitube) for 15 min. Lysates were diluted 1:2 in ChIP dilution buffer to neutralise the SDS and pre-cleared with Protein G Dynabeads (ThermoFisher Scientific). Immunoprecipitation was performed overnight at 4°C using 5mg antibody per 3 x 10^5^ cells. 10% of input was frozen for input controls. Beads were added for 2 hr and washed sequentially with low-salt, high-salt, LiCl, and Tris buffers.

Bead-bound chromatin was tagmented for 10 min at 37°C (1,400 rpm) using loaded Tagmentase bound with sequencing adaptors (Diagenode). DNA was eluted, reverse cross-linked overnight at 65°C with Proteinase-K, and purified by phenol-chloroform extraction. Libraries were amplified using unique dual indices (Diagenode) and Kapa HiFi PCR Master Mix (Roche), purified with Kapa Pure beads, and assessed on an Agilent Bioanalyzer (High sensitivity DNA chip). Libraries were quantified by qPCR (NEBNext Library Quant Kit) and sequenced on a NovaSeq 6000 (Illumina) to a depth of 50-100 million reads per sample. Data were processed using the nf-core chipseq v1.2.1 pipeline^30^ with default parameters.

### Omni-ATAC

ATAC-seq was performed as described by Corces *et al*^31^. Briefly, 5 x 10^5^ cells with >90% viability were lysed in ice cold RSB buffer containing 0.1% NP40, 0.1% Tween 20 and 0.1% Digitonin for 3 mins. Lysis was quenched with 1ml cold RSB buffer, and nuclei were pelleted and resuspended in transposition mix containing 100nM loaded Tn5 transposase (Diagenode). Tagmentation was performed at 37°C for 30 mins with shaking (1,000 rpm). Eluted DNA was amplified using UDI adaptors (Set I-II, Diagenode) and Kapa HiFi PCR master mix (Roche) for 7-10 cycles. Library quality was assessed using a Bioanalyzer DNA kit (Agilent Technologies) and quantified by qPCR (NEB Library Quant Kit) on a QuantStudio7 Real Time PCR (Thermo Fisher Scientific). Libraries were sequenced as 50bp paired-end reads (70-100 million reads per sample) on a NovaSeq 6000 (Illumina) using an S4 flow cell. Data were processed using the nf-core atacseq v1.2.1 pipeline^32^ with default parameters.

### RNA extraction and sequencing

Total RNA was extracted from 1 x 10^6^ cells using the RNeasy Mini Kit (Qiagen). RNA quantity and integrity were assessed by Qubit 3.0 fluorometry (Life Technologies) and Agilent Bioanalyser; all the samples had RNA integrity numbers (RIN) >9.3. Ribosomal RNA-depleted libraries were prepared with unique barcodes and sequenced as 100 bp paired-end reads on a NovaSeq 6000 (Illumina). Data were processed using RNAflow^33^ v1.4.1. Differential expression analysis was performed using DESeq2, comparing each cancer cell line to the normal colon cell line (HCEC-1CT). *P*-values were adjusted using Benjamini-Hochberg procedure, with *FDR* < 0.05 considered significant.

### gRNA-library design

Candidate enhancer regions were identified by integrating ATAC-seq, H3K27ac and H3K4me1 ChIP-seq peaks from CRC cell lines C32, CL11, HT29, SW403, SW480, and SW948. Peak BED files were intersected using Intervene^34^, and regions present in ≥5 cell lines were retained. Guide RNAs (gRNA) were designed as 20bp oligos using FlashFry^35^ v1.12, selecting up to three gRNAs per enhancer. In addition to the 35,139 gRNAs mapping to 12,117 enhancer regions, we included 100 non-targeting controls (NTC): 50 non-targeting scrambled-sequence spacers and 11 protospacers targeting gene-devoid genomic regions as per Gasperini *et al*.^4^, and 39 NTCs as per Horlbeck *et al*.^36^

### gRNA-library cloning and transduction

Guide oligonucleotides containing vector-flanking sequences (Twist Bioscience) were synthesised as 5’-atcttgtggaaaggacgaaacaccG-N_20_-gtttaagagctatgctggaaacagcatagcaagt-3’. gRNA cloning was performed as described by Gasperini *et al*^4^. Oligonucleotides were PCR-amplified (12 cycles) using Q5 High-Fidelity Master Mix (NEB), purified (DNA Clean & Concentrator-5, Zymo Research), and inserted into the CROP-seq-opti vector (Addgene #106280) by Gibson assembly (NEBuilder HiFi, NEB). Following assembly, 100ng plasmid was transfected into NEB High Efficiency Stable Competent *E. coli* cells (1.8 kV, Eppendorf Eporator) and plated on carbenicillin (500µg/ml) agar. 2 x 10^6^ colonies were harvested, and plasmid DNA purified using ZymoPURE Maxi kits (Zymo Research).

Lenti-viral transduction of cells to incorporate the CRISPR-dCas9 gene was performed in HEK239T cells as described by Gordon *et al*^37^. Lentivirus particles were generated in HEK239T cells. Per T175 flask, 10µg pHR-UCOE-SFFV-dCas9-mCherry-ZIM3-KRAB (Addgene #154473), 6.5µg psPAX2 (Addgene #12260), and 3.5µg pMD2.G (Addgene #12259) were diluted in 2ml of Opti-MEM (Gibco) and 40µl of TurboFect (Thermo Scientific). Viral particles were concentrated (Lenti-X Concentrator, Takara Bio) and MOI determined using Lenti-X GoStix Plus (Takara Bio). HT29 cells (∼8 x 10^6^) were transduced in the presence of polybrene (8µg/ml) and mCherry-positive cells isolated by flow cytometry (Beckton Dickinson Symphony S6 Cell Sorter).

Lentivirus containing the gRNA library (CROP-seq-opti vector, Addgene #106280) was produced similarly. At least 5 x 10^6^ HT29-dCas9 cells were transduced to integrate the lentiviral particles containing the CROP-seq-opti vector at an MOI ≥ 80. After 48 hrs cells were selected with puromycin (2ug/mL). Cells were harvested 10 days post-transduction for single-cell RNA sequencing.

### Sequencing of scRNA-seq libraries

Single-cell transcriptome capture was performed using the Parse WT-Mega v1 kit (Parse Biosciences), according to manufacturer’s instructions. To enrich for gRNA sequences, a three step semi-nested PCR was performed on barcoded cDNA as described by Gasperini *et al*^4^. Libraries were pooled and divided into 12 sub-libraries, and sequenced on a NovaSeq 6000 (Illumina) using an S4 flow cell with cycle configuration 120-6-86. Sequencing data were processed using the split-pipe pipeline v1.7.1 (Parse Biosciences). Reads were aligned to GRCh38.p13, with gene annotations from Ensembl v105. The reference genome was modified to include gRNA sequences as annotated “genes” to enable read assignment (**Table 2**). The resultant expression matrix was partitioned into gene and gRNA count matrices for downstream analyses.

**Table 2:** Table of primers. Shown are the primers used for cloning and library preparation, as well as the sgRNA “gene” definition used for guide detection.

### Differential gene expression

Differential expression analysis was performed using the sceptre^11^ package v0.10.3, which applies a resampling framework optimised for Perturb-seq experiments. The standard SCEPTRE workflow was implemented, including gRNA assignment using a mixture model accounting for cell-specific covariates, cell- and pair-level quality control, calibration checks, and discovery analysis (**Fig. 3**).

Cells with extreme numbers of detected genes or UMIs (outside the 1st-99th percentiles) or ≥20% mitochondrial reads were excluded. Enhancer-gene pairs were only retained if ≥7 perturbed and ≥7 control cells exhibited non-zero expression of the gene. After QC, 308,796 gRNA-gene associations within 2Mb of a transcription start site (TSS) were tested (**Fig. 3**).

Negative control region-gene pairs were generated by grouping three non-targeting gRNAs and analysed using the same QC criteria.

Associations were calculated tested using negative binomial regression with conditional resampling, including the following covariates: expressed genes, gene read depth, number of expressed gRNA, gRNA read depth, mitochondrial read fraction, and sequencing batch. A left-tailed test was applied to detect decreases in expression. *P*-values were adjusted using Benjamini-Hochberg procedure, and associations with at *FDR* < 0.1 considered significant.

### Micro-C and ABC analysis

Micro-C libraries were generated for HT29, CL11, SW403, SW480, and SW948 as described previously^38,39^. Briefly, cells (1 x 10^6^ cells/ml) were crosslinked with DSG (3mM) followed by 1% formaldehyde, digested with MNase (Worthington), end-repaired and biotin-labelled, proximity-ligated, and reverse cross-linked. Size selected fragments (250-400bp) were purified, captured with streptavidin beads, adaptor-ligated (NEB), PCR-amplified, and quality assessed prior to sequencing. Libraries were sequenced on a NovaSeq 6000 (Illumina) to ≥300 million 100bp paired-end reads. Quality control was performed using JuicerTools^40^, required >90% cis contacts and 60-70% short range cis interactions

Interaction maps were generated using nf-distiller^41^ v0.3.4, and normalised using Raichu^42^ v1.1 at 1kb, 5kb, and 10kb using the same parameters as Wang *et al*. Significant contacts were identified using pyHICCUPS v0.3.9. TADs and compartments were called using cooltools^43^ v0.5.4 at 30kb and 100kb windows respectively. Compartments were defined by eigenvector decomposition (10 kb bins) and phased by GC content. Eigenvector values > 0 were defined as active compartments.

Enhancer-gene links were predicted using the ABC model^15,16^ v1.1.2 integrating ATAC-seq, H3K27ac ChIP-seq, Micro-C, and RNA-seq data. Analyses followed Nasser *et al*.^16^, restricting candidate enhancers to those tested in the Perturb-seq analysis and genes within 2Mb.

### Transcription Factor binding

Transcription factors (TF) occupancy within candidate enhancers was predicted using TOBIAS^44^ v0.14.0 with the ATAC-seq data and the JASPAR 2022 core non-redundant motif database^45^, filtered to motifs found in humans. TFs were retained if the TOBIAS score was > 1 and binding was predicted in ≥10% of candidate enhancers.

### Functional annotation of enhancer-gene relationships

Random forest classification was performed to identify features associated with functional enhancer-gene pairs. Positives were defined as SCEPTRE associations with FDR < 0.1; negatives as those with an FDR > 0.1 and log_2_-fold-change between -0.05 and 0.05. Features included genomic context (intronic, upstream), ABC score, 3D contact metrics, TAD colocalization, compartment status, GWAS overlap, and quantile-normalized ATAC, CTCF, H3K27ac, H3K4me1 and H3K4me3 signals at enhancers and promoters^15^. Gene-level features included housekeeping status, transcript length, and strand. To address class imbalance, negatives were down-sampled to 10,000 and balanced class weights applied. Feature importance was assessed by mean decrease in impurity using scikit-learn^46^ v1.2.1.

The same dataset was evaluated using the model framework of Gschwind *et al*^17^. ENCODE-predictions were obtained for HT29 (ENCODE annotation set ENCSR602YDS), GM12878 (ENCSR252PIL), and K562 (ENCSR328LMT).

### Gene evidence

Gene set analyses were performed using oncoEnrichR^47^ v1.4.2.1, which integrates cancer associations, drug associations, synthetic lethality, gene fitness, and protein-protein interaction data.

Regulatory interaction data were obtained from the OmniPath^48^ and DoRothEA^49^, which curate TF-target interactions supported by literature annotation, ChIP-seq evidence, TF binding site motifs, and expression-based inference. Regulatory interactions were further inferred using pan-cancer expression data from TCGA.

Gene essentiality and cell viability data were obtained from The Cancer Dependency Map (DepMap; 2020_Q2 release), comprising genome-scale CRISPR/Cas9 knockout screens across 912 cancer cell lines^50^. Analyses were restricted to colorectal carcinoma cell lines (37 primary, 14 metastatic). Putative therapeutic targets were prioritised using DepMap Project Score (20210311 release)^50,51^, which integrates CRISPR fitness effects with genomic biomarker and tumour prevalence data. Target priority scores range from 0-100, with higher scores indicating greater prioritisation. A threshold score ≥40 was applied, based on benchmarking against approved or preclinical oncology targets^50^.

## DECLARATIONS

### Availability of data and materials

The datasets supporting the conclusions of this article are available in the European Genome-phenome Archive (EGA; https://ega-archive.org/) repository under the following accessions: EGAD50000000375 (Perturb-seq), EGAD50000000294 (Micro-C), EGAD50000000295 (ChIP-seq, all marks), EGAD50000000296 (ATAC-seq), EGAD50000000297 (RNA-seq).

### Authors’ contributions

JS, PJL and RSH conceived and designed the study; JS, JV and MM performed experiments and data generation; PJL performed computational and statistical analysis, and data integration and interpretation; TB and EK provided sceptre support; PJL and RSH drafted the manuscript. All authors have read and approved the final manuscript.

## Acknowledgements

We thank Andrew Everall and Andrea Gunnell for statistical and technical input.

